# Alpha-band frequency and temporal windows in perception: A review and living meta-analysis of 27 experiments (and counting)

**DOI:** 10.1101/2023.06.03.543590

**Authors:** Jason Samaha, Vincenzo Romei

**Affiliations:** Department of Psychology, University of California, Santa Cruz, USA; Centro studi e ricerche in Neuroscienze Cognitive, Dipartimento di Psicologia, Alma Mater Studiorum – Università di Bologna, Campus di Cesena, Cesena, 47521, Italy; Facultad de Lenguas y Educación, Universidad Antonio de Nebrija, Madrid, 28015, Spain

## Abstract

Temporal windows in perception refer to windows of time within which distinct stimuli interact to influence perception. A simple example is two temporally proximal stimuli fusing into a single percept. It has long been hypothesized that the human alpha rhythm (an 8-13 Hz neural oscillation maximal over posterior cortex) is linked to temporal windows, with higher frequencies corresponding to shorter windows and finer-grained temporal resolution. This hypothesis has garnered support from studies demonstrating a correlation between individual differences in alpha frequency (IAF) and behavioral measures of temporal processing. However, non-significant effects have also been reported. Here, we review and meta-analyze 27 experiments correlating IAF with measures of visual and audio-visual temporal processing. Our results estimate the true correlation in the population to be between 0.39 to 0.53, a medium-to-large effect. The effect held when considering visual or audio-visual experiments separately, when examining different IAF estimation protocols (i.e., eyes-open and eyes-closed), and when using analysis choices that favor a null result. Our review shows that 1) effects have been internally and independently replicated 2) several positive effects are based on larger sample sizes than the null effects and 3) many reported null effects are actually in the direction predicted by the hypothesis. A free interactive web-app was developed to allow users to replicate our meta-analysis and change or update the study selection at will, making this a “living” meta-analysis (https://randfxmeta.streamlit.app). We discuss possible factors underlying null reports, design recommendations, and open questions for future research

## Background

Our brains routinely integrate perceptual information within and across sensory modalities. In many cases, temporal coincidence is a key factor in determining whether two inputs become integrated or segregated. For instance, in the well-known sound-induced flash illusion (SIFI; see Figure 1), a single flash can be perceived as two separate flashes if accompanied by two beeps, with the second beep occurring within a time window of ∼100 ms of the flash (Shams et al., 2000, 2002), suggesting audio-visual interactions occurring within a critical temporal window. There is no single integration window for all types of stimulus features and sensory modalities. Indeed, many perceptual phenomena, particularly motion-related ones, imply a range of integration windows spanning up to hundreds of milliseconds (Herzog et al., 2020) and even seconds (Liu et al., 2019). However, integration windows for many low level features fall within 100 ms. For instance the time needed to discriminate one from two spatially overlapping light flashes is on the order of 40-60 ms in adults (Exner, 1875; Pearson & Tong, 1968), as is the time needed to perceive synchronicity between two visual (Milton & Pleydell-Pearce, 2016; Varela et al., 1981) or an auditory and visual event (Kristofferson, 1967b). Many visual masking phenomena also occur with maximum strength at intervals between 50 and 100 ms (Breitmeyer & Ganz, 1976; Ro, 2019), and a number of space/time based visual illusions, such as the flash-lag and Fröhlich effect are consistent with perceptual displacements on the order of 30-80 ms (Chakravarthi & VanRullen, 2012; Morrow & Samaha, 2022; Schneider, 2018).

**Figure 1.**
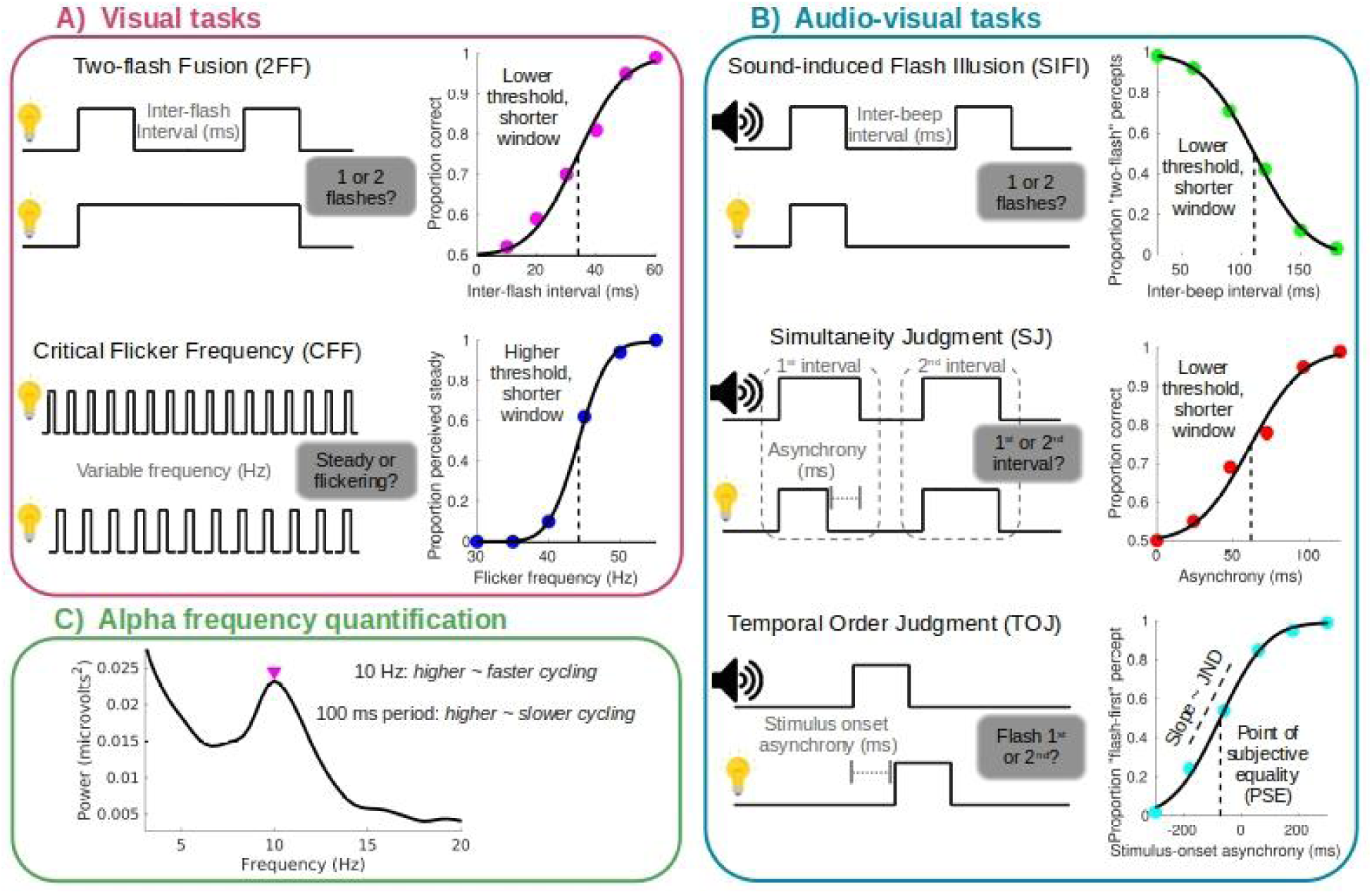
Common behavioral measures of temporal binding windows and IAF measurement approaches. **A)** Common visual tasks. In the two-flash fusion task (2FF) subjects report whether they perceived a single flash or two separate flashes, with the critical manipulation being the duration between flashes, or inter-flash interval. In some studies, only two-flash trials are presented (making it a detection task potentially susceptible to bias), whereas other implementations (as shown here) present duration-matched single-flash trials as well (making it a discrimination task that affords the quantification of both sensitivity and bias). Typically the middle point of the psychometric function is taken as a threshold which indicates the inter-flash interval at which two-flashes are reliably discriminated from one. A lower 2FF threshold (in ms) indicates a shorter temporal binding window (or higher temporal resolution). The critical flicker frequency (CFF) has long been used in research and clinical settings as a measure of temporal resolution. To measure CFF, a flickering light of varying frequency is presented and the threshold frequency at which a subject reliably perceives the flickering light as steady illumination is recorded. Here, a higher threshold (in Hz) corresponds to a shorter binding window (or higher resolution). **B)** Common audio-visual tasks. In the sound-induced flash illusion (SIFI) subjects report whether they perceived one or two flashes. Often, a single flash accompanied by two beeps will induce the illusion of two flashes. This occurs with higher probability at shorter inter-beep intervals. In the SIFI, a lower threshold (in ms) indicates a shorter window within which the second beep induces an illusory second flash percept. Simultaneity judgment tasks (SJ) require the subject to indicate whether two stimuli appear to offset (or onset) at the same time or not. The bias-free, 2-interval forced-choice (2IFC) version of this task used by Kristofferson (1967b, 1967a) is depicted here. Subjects judged which of two intervals contained an audio-visual pair with an asynchronous offset. The degree of asynchrony (in ms) required to reliably choose the correct interval corresponds to the threshold, with lower values indicating shorter audio-visual binding windows. In the temporal order judgment task (TOJ) an audio-visual pair is presented with a varying degree of stimulus onset asynchrony (SOA) and subjects judge which stimulus was presented first (sometimes phrased as whether the flash was presented first or second). A typical psychometric function from this task can be quantified in terms of its slope, which is related to the just noticeable difference (JND), and is steeper when the subject has better temporal resolution (shorter binding window). Here, the threshold defines the point of subjective equality (PSE) - the SOA at which the audio and visual stimuli appear simultaneous. **C)** A typical EEG power spectrum recorded from electrode Oz (first author’s data). IAF is commonly defined as the frequency with max power within the alpha range (i.e., 7-14 Hz) and can be obtained from eyes-open (as in this example) or eyes-closed recordings. IAF may also be derived from a model fit to the power spectrum (such as fitting a Gaussian curve to the whole alpha “bump”) or via a “center-of-gravity” approach (which amounts to a amplitude-weighted average of the entire alpha range). The peak is sometimes reported in Hz, with higher values corresponding to “faster” cycling, or in terms of the oscillation’s period in ms (1000*(1/Hz)) with higher values corresponding to “slower” cycling. Importantly, we sign-corrected each correlation in the meta-analysis to account for the IAF metric (Hz or ms) and the direction of the behavioral metric (longer/shorter window size).

The coincidence of many temporal integration phenomena occurring within 100 ms and the average period of the human alpha rhythm (∼100 ms or 10 Hz) has long motivated the hypothesis that the two are related (Harter, 1967; VanRullen, 2016), with a key prediction being that the *frequency* of alpha activity (which differs across and within individuals) is related to the duration of temporal windows in one’s perception. This hypothesis garnered empirical support in the second half of the 20th century from a number of experiments (Coffin, 1977; Coffin & Ganz, 1977; Kristofferson, 1967b; Murphree, 1954; Varela et al., 1981) and in recent years, growing interest in the topic has led to many studies examining the link between alpha frequency and temporal properties of perception. In this paper, we review those studies, provide a meta-analysis of published studies in order to estimate the presence and strength of any such relationship, and attempt to address recent null findings (Buergers & Noppeney, 2022). Given the likely continued growth of this research area, we have programmed a free, open-source, interactive web app that replicates our meta-analysis (https://randfxmeta.streamlit.app). This allows any user to easily update meta-analytic estimates as new studies become available, to see what the results would look like under different study inclusion/exclusion conditions, or to conduct their own meta-analysis on any correlational data.

Our focus is on reviewing empirical work that correlates individual differences in alpha-band frequency (IAF) with individual differences in temporal aspects of visual and audio-visual perception, as these are the most highly studied modalities. Although different mechanistic explanations have been proposed to underlie the link between alpha and temporal windows (Dou et al., 2022; Morrow et al., 2023; VanRullen, 2016; VanRullen & Koch, 2003), we put aside the question of mechanism for now to focus on the strength of existing empirical evidence motivating such mechanistic explanation in the first place. To facilitate comparison across different studies, we focused just on experiments that correlate between-subject variation in IAF with a behavioral index of temporal processing (see Figure 1). There is also relevant evidence from experiments that examined within-subject variation in alpha frequency which we touch upon in the discussion, but there are not enough of such studies for meta-analysis. Moreover, the effects of alpha-phase (Ruzzoli et al., 2019; VanRullen, 2016) and power (Samaha et al., 2020) on perception have recently been reviewed elsewhere and are not discussed here. We begin by reviewing evidence from the visual domain relating alpha frequency and temporal properties of perception.

## Alpha-band frequency and the temporal properties of visual perception

In a pioneering yet not widely cited study, Coffin and Ganz were among the first to test for a link between alpha frequency and temporal binding within the visual domain (Coffin & Ganz, 1977). Their studies used a two-flash fusion (2FF; Figure 1A) task to measure the probability that observers perceived two brief visual flashes separated by a threshold-level inter-stimulus interval (ISI; usually around 40 ms) as a single flash. In 9 observers they found that trials where two flashes were reported instead of one were accompanied by higher prestimulus peak-alpha frequencies measured over the occipital pole, suggesting that visual alpha happened to be cycling at a faster rate when the two stimuli were successfully individuated. A follow up experiment in 13 subjects used a signal-detection theory (SDT) framework to determine if alpha frequency shifts were associated with a change in criterion (tendency to report one or two stimuli independent of the actual stimulus) or sensitivity (enhanced discrimination of one versus two flashes). Using a 12 ms ISI (which was virtually always perceived as a single flash) versus a threshold ISI (perceived as two flashes ∼ 75% of the time), they determined that higher alpha frequencies lead to greater hits (reporting “two-flashes” at threshold ISI), without changing false alarms (reporting two-flashes at the 12 ms ISI), linking alpha frequency to the temporal sensitivity of visual processing within subjects.

Decades later, a similar effect was shown in an experiment by Samaha & Postle (2015), who expanded on the 2FF task design and EEG analysis approach in several ways. First, Samaha & Postle (2015) tested a range of ISIs in each observer and, importantly, included an equal number of one-flash and two-flash trials in order to measure the accuracy of observers’ ability to discriminate one from two flashes across their psychometric functions. An important consideration in this design was controlling for the possible confound that observers base their flash perception judgments, in part, on the overall duration of the stimulus (since, all else being equal, two flashes separated by a long ISI will be a longer stimulus event than two flashes separated by a short ISI or a single flash). Samaha & Postle (2015) accomplished this by varying the length of each one-flash stimulus to match that of each possible two-flash sequence, thereby removing stimulus duration as a confounding cue. The study found that, within-subjects, trials with faster pre-stimulus alpha frequency predicted two-flash discrimination accuracy (akin to Coffin & Ganz) but also that individual differences in both eyes-closed and pre-stimulus IAF negatively correlated with 2FF thresholds (the ISI producing 75% correct discriminations). Topographical analysis revealed medium-high correlations (*r* around -0.5) between IAF and 2FF threshold across bilateral occipital electrodes, an effect that held also when controlling for participant age and fatigue levels. The direction of the effect indicates that higher alpha frequency corresponds to finer-grained temporal resolution (or shorter temporal binding window), but how consistent is this pattern of results?

We found 27 experiments that have examined the correlation between IAF and temporal properties of visual and audio-visual perception (summarized in Table 1). The majority of studies (14) have looked at purely visual phenomena. For instance, in one of the earlier studies Sokoliuk & VanRullen (2013) found that IAF strongly correlated with the frequency of illusory flicker perception in the flickering wheel illusion (r ∼ 0.8), a phenomenon whereby a static, high contrast, high spatial frequency “wheel” is seen to flicker when viewed peripherally. More obviously temporal phenomena have also been linked to alpha frequency such as the illusion of motion-induced spatial conflict (also known as the “fluttering heart illusion”). In this illusion, a moving border defined only by color contrast (and not by luminance) is seen to jitter or flutter as it moves, perhaps as a result of timing differences between motion and shape processing channels (Arnold & Johnston, 2003). Recently, Minami & Amano (2017) showed that the precise frequency of the perceived jitter is strongly correlated with occipital IAF (r ∼ 0.8). The authors also provided causal evidence by modulating IAF with alternating current stimulation (tACS) above or below each individual’s peak alpha, which could speed up or slow down the rate of illusory flicker.

**Table 1.**
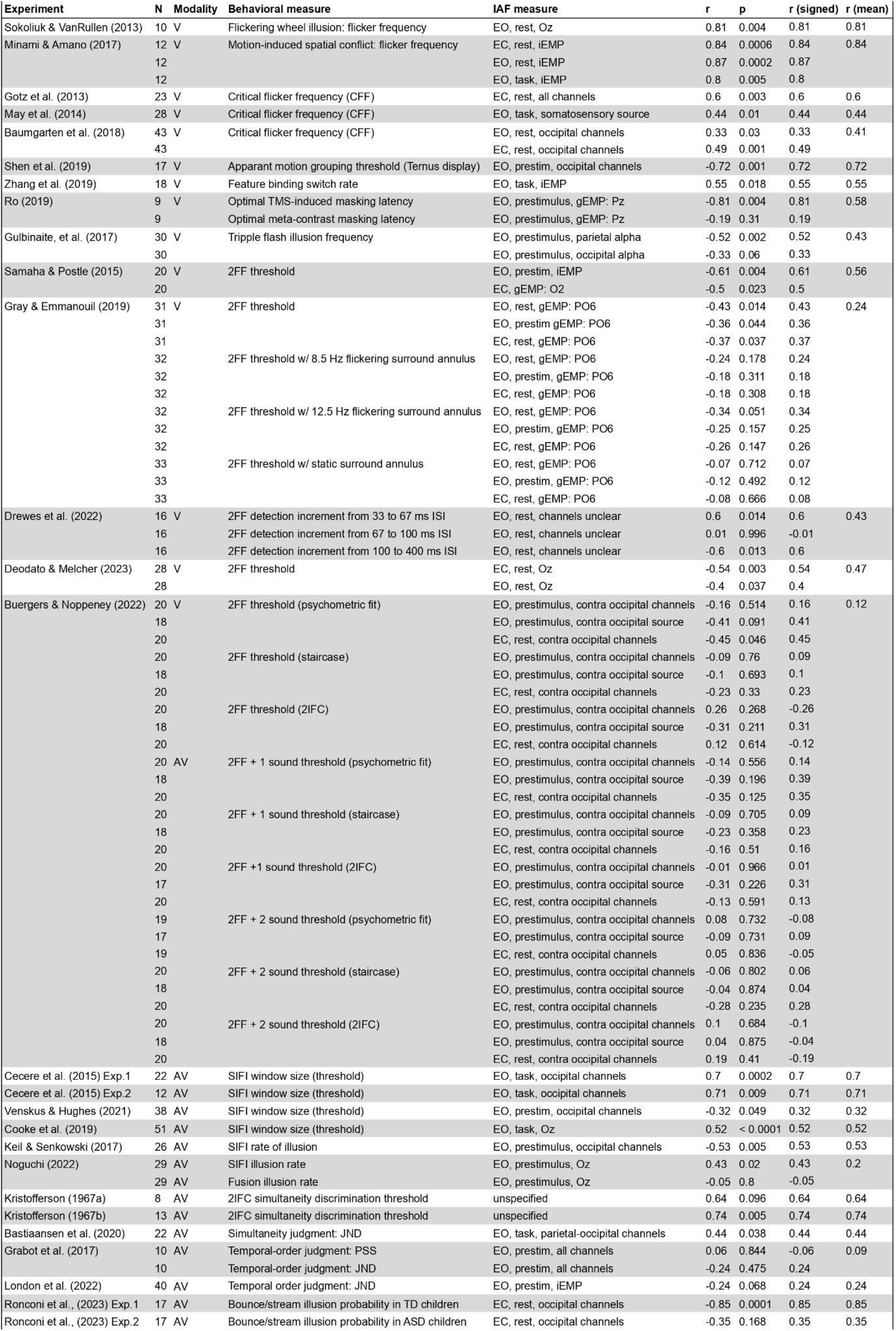
Abbreviations: group (g); individual (i); electrode w/ max power (EMP); two-flash fusion (2FF); visual (V); audio-visual (AV); just noticeable difference (JND); point of subjective simultaneity (PSS); sound-induced flash illusion (SIFI); typically developing (TD); autism spectrum disorder (ASD)

In addition to illusory flicker phenomena, temporal binding has also been investigated in cases such as the Ternus display, whereby a group of moving dots can either be perceived as individually moving or moving a group depending on the ISI between frames. Shen et al. (2019) found a strong correlation (r ∼ 0.7) between the ISI at which perception switches between the two motion interpretations and IAF. They verified this link not only in EEG data using individual differences, but also in intracranial recordings from visual areas using trial-to-trial variation in alpha frequency to predict perceptual interpretations (Shen et al., 2019). Fluctuations in feature binding over time have also been found to relate to IAF. Using a display which induced a misbinding between color and moving dots, Zhang et al. (2019) found that higher IAF predicted more frequent switches between the correct and incorrect feature binding.

Notably, the original result from (Samaha & Postle, 2015) was independently replicated in a larger sample by Gray & Emmanouil (2020) who used identical stimulus parameters in a 2FF design and also by several other groups who used similar stimuli (Deodato & Melcher, 2023; Drewes et al., 2022). However, when Gray & Emmanouil (2020) measured 2FF thresholds under new conditions that included a high-contrast flickering annulus surrounding the target flash stimuli (in an attempt to entrain alpha), they no longer observed a significant correlation between IAF and 2FF (although the correlations were all still in the same direction, see Table 1). This is perhaps not so surprising since the presence of the annulus, even when static, significantly altered 2FF thresholds, suggesting some low-level interactions between the target flashes and the annulus.

A straightforward and clinically-relevant measure of the temporal resolution of vision is the critical flicker frequency (CFF; Figure 1A), which refers to the lowest frequency of flickering light required to achieve a non-flickering percept (i.e., a percept of constant illumination). CFF has predictive utility as a clinical biomarker, in particular, for classifying the severity of haptic encephalography (HE). Notably, HE is also associated with a reduction of alpha frequency, which motivated the authors of three separate papers to look at the link between IAF and CFF in relatively large sample sizes that comprise both HE patients and controls. These three studies all report positive correlations (ranging between 0.33 and 0.6), indicating that higher IAF is associated with higher visual temporal resolution (Baumgarten et al., 2018; Götz et al., 2013; May et al., 2014). Although these studies typically pool data across clinical and control groups, this provides measures that vary more across both IAF and CFF than a typical college sample, giving an opportunity to study the relation between the two variables across a wider measurement range. Although one cannot infer causality from these results, the fact that a condition associated with lower IAF is also associated with lower temporal resolution could be seen as evidence in favor of the hypothesis.

Table 1 summarizes all the studies we found, including sample size information, task information, IAF computation details, and whether the effects are in the direction suggested by the theory (since different behavioral measurements or representations of alpha frequency as being in period length in ms or in Hz can arbitrarily flip the sign of the correlation; see Figure 1). In later sections we provide a more complete meta-analytic description of these studies, but for now we note that the large majority of studies in the visual domain find support for the idea that IAF is related to temporal properties of perception and that, where binding window durations can be directly estimated from the behavioral measure, higher IAF corresponds to higher temporal resolution. A notable exception is the result from Buergers & Noppeney (2022) who tested 20 participants across multiple days using 2FF tasks that varied in the number of sounds simultaneously presented from zero sounds (visual only), one sound, and two sounds. They additionally varied the type of task used to measure binding window thresholds by using the method of constant stimuli in a 2FF discrimination task which could be fit with a psychometric function, a staircase procedure, and a two-interval forced choice task (2IFC). The results failed to replicate the within-subject effect of alpha frequency on 2FF discrimination and also the between-subject correlation with IAF and 2FF thresholds across the various task and sound manipulations (see Table 1). We will later discuss some possible methodological reasons for the null result, but for now we note that many of the between-subject correlations reported are actually in the expected direction but are not statistically significant. Moreover the predicted negative correlation between IAF and 2FF threshold in the visual-only condition was actually significant (two-tailed) when eyes-closed IAF was derived from electrodes contralateral to the stimulus location and when pre-stimulus IAF was derived from source estimated activity in contralateral occipital cortex (one-tailed), signals which perhaps better capture alpha activity in the neural population relevant for processing the lateralized visual stimulus (Table 1).

## Alpha-band frequency and multi-sensory perception

The relevance of temporal binding becomes even more clear when it comes to combining information across the senses. An issue arises for the brain due to different sensory stimuli having different inherent temporal properties. For example, light travels at a much faster speed than sound, a well-known phenomenon one might witness during a thunderstorm. In addition to the differential physical propagation of audio and visual stimuli, there are also well-known neural processing differences. The visual system generally has higher spatial precision than the auditory system but lower temporal precision. This means that perception of simultaneity between two audio-visual stimuli must have a certain degree of flexibility such that we can perceive two stimuli as simultaneous even when they arrive and are processed with different latencies, i.e., as long as they are within a certain temporal window. This window may have been adapted to accommodate for the physical properties of the stimuli we receive, which, depending on the distance of the source, may reach our senses at different times. Across different tasks, and especially so for simple stimuli, the exposure to multisensory stimulation with variable onset differences may lead to a certain level of tolerance for simultaneity with a temporal window measured to be around 100ms. Variability on the width of this temporal binding window can be observed both for intra and inter-individual measurements.

Some of the most common tasks used to quantify the width of the audio-visual temporal binding window are the simultaneity judgment (SJ) task, the temporal order judgment (TOJ) task, and the sound-induced flash illusion (SIFI) described above (see Figure 1B). Although these tasks seem, in concept, to tap into the same underlying binding process, estimates across tasks are sometimes found to correlate but sometimes not (for discussion see Kostaki & Vatakis, 2018). One important factor may be that different temporal sensitivities across the senses could lead to different temporal binding window estimates depending on the leading sense. For example, whether the auditory or visual stimulus leads or lags.

In two landmark studies which were perhaps the first to directly test the link between temporal binding and alpha frequency, Kristofferson (1967b, 1967a) used a SJ task to measure the threshold interval at which subjects could discriminate between a simultaneously offsetting light-sound pair and a non-simultaneously offsetting stimulus where the light terminated before the sound. Importantly, judgments were made in a 2IFC design whereby subjects were presented with both a simultaneous and non-simultaneous stimulus in separate intervals of each trial and had to decide which interval contained the non-simultaneous stimulus. SJ thresholds estimated this way cannot be biased by an overall propensity to judge stimuli as simultaneous or not and thus better reflect participants sensitivity thresholds for perceiving audio-visual simultaneity. In a first study with 8 subjects, discrimination thresholds were found to correlate at 0.64 with alpha period, revealing that faster alpha predicts a lower SJ threshold (Kristofferson, 1967a). A subsequent study with 13 participants replicated the effect and found a significant correlation of 0.74 (Kristofferson, 1967b).

Decades later Cecere et al. (2015) demonstrated that the temporal binding window in the SIFI can be predicted by one’s IAF, using both a correlational and a causal approach. Their study found that, in 22 subjects, individuals with lower frequency occipital IAF had elongated windows within which the second sound caused the illusory perception of a second flash (r = 0.70), indicating audio-visual interactions occur over a longer time window the lower one’s IAF. A follow-up study using tACS found that the SIFI window could be expanded or contracted by stimulating below or above IAF, respectively.

As shown in Table 1, the correlation between IAF and the SIFI has been replicated by the same group in a larger sample (n=51; Cooke et al., 2019) and by three other groups (Keil & Senkowski, 2017; Noguchi, 2022; Venskus & Hughes, 2021). Of note, Noguchi (2022) (see also Noguchi, 2023) used SDT analysis to show that the link between SIFI and IAF is not only present when looking at the frequency with which an individual experiences the illusion, but is specifically linked to changes flash-discrimination sensitivity (and not criterion). Interestingly Noguchi (2022) also found that IAF was uncorrelated with the sound-induced ‘fusion’ illusion, whereby two flashes presented with a single beep can sometimes cause fusion of the flashes and a percept of a single flash. Instead, in the fusion case, individual beta frequency was found to correlate with the illusion frequency, suggesting a dissociation between beta and alpha frequencies in the fusion or ‘fission’ (i.e., the classic SIFI) of audio-visual stimuli.

Beyond the SIFI, recent work has also replicated links between IAF and audiovisual binding as measured in the SJ task, akin to the early work by Kristofferson (1967b, 1967a). A study by Bastiaansen et al. (2020) found that, in 22 subjects, higher IAF predicted more sensitive audio-visual simultaneity perception (lower just-noticeable differences; JND) in a task using visual-leading stimuli. Experiments using the TOJ task have found less consistent effects, however. A study with 10 subjects found no significant correlation between JND and IAF (although the effect was in the expected direction; (Grabot et al., 2017)) and a larger study of 40 subjects found only a trending association between IAF and JND in the expected direction (r = -0.24, p = 0.06; (London et al., 2022)). Exploring the effect of the leading sense could be interesting since, in the TOJ task, the order can be judged for both visual-leading and auditory-leading stimuli. Lastly, a recent paper examined temporal windows of audiovisual integration in the so-called “bounce/stream” illusion. In this effect, two dots move towards each other on a screen, coincide, and then continue along their trajectories. This often appears as two dots “streaming” past each other, but if a “bounce-like” sound occurs around the time of dot coincidence, the dots tend to appear as though they bounced off of one another. Ronconi et al. (2023) found a strong correlation between the probability of bounce percepts and IAF in 17 typically developing (TD) children (r = -0.85), indicating that bounce percepts decreased as IAF increased, suggesting smaller windows the higher one’s IAF. Interestingly, a second experiment with 17 children with autism spectrum disorder (ASD) found a much weaker relationship (r = -0.35).

As noted in the section on visual effects, the experiments of Buergers & Noppeney (2022) also failed to replicate some of the audio-visual findings reviewed above, in particular regarding the SIFI. A trend notable in Table 1, however, is that even many of the reported null effects were in the direction expected under the hypothesis. Therefore, to get a better overall estimate of the presence and strength of the proposed relationship between IAF and temporal windows, we conducted a meta-analysis of all 27 experiments reported in Table 1.

## Meta-analytic methods and results

Table 1 summarizes all 27 experiments we found that correlated IAF with a behavioral measure of the temporal properties of visual or audio-visual perception. We included data on the correlation r and p-value, the sample size, sensory modality, the behavioral measure, and the quantification of IAF. We also included a column interpreting the sign of the correlation in terms of the prediction that higher IAF predicts shorter binding windows/more frequency updating. We discovered studies using a backwards and forward citation search of several highly-cited classical and modern studies in this field (Cecere et al., 2015; Kristofferson, 1967b; Samaha & Postle, 2015), and by searching PubMed with the following keyword combinations: “alpha frequency” [AND] “binding”, “alpha frequency” [AND] “temporal resolution”, “alpha frequency” [AND] “multisensory integration”, “alpha frequency” [AND] “sampling”, “alpha frequency” [AND] “flicker”. Although records were not kept of every study screened, we note that readers can add or remove selected studies and observe the effect on meta-analytic outcomes using the free, interactive web app that we provide (https://randfxmeta.streamlit.app).

Many studies reported a correlation using different derivations of IAF (e.g., using several different electrode groups or using eyes-opened and eyes-closed measurements). As a result, the total number of correlations we compiled was 75, even though there are only 27 unique experimental samples. We therefore also included a column that tracks the mean correlation (called r(mean) in the table) for each unique experiment, averaging over the different IAF derivations or study conditions, which should be tapping into the same underlying constructs. Importantly, we also computed the “sign-corrected” r-value by flipping the sign of the correlation in cases where a negative correlation was actually in the direction of the theory (for example when higher IAF predicts a lower 2FF threshold, see Figure 1). This corrected r-value (called r(signed) in the table) is positive when a correlation supports the theory and negative if it goes against the theory. As noted in Table 1, the direction of two correlations we found were unclear to us from a theoretical perspective, so we counted those as going against the theory in order to reduce bias towards a positive result. We use this sign-corrected correlation in all subsequent analyses. The table is sorted by modality (visual or audio-visual) and then by behavioral measure, to facilitate comparison across studies using similar approaches. Data in this table, along with matlab code used for the meta-analysis described below can be found at https://osf.io/sgkum/.

We used a random-effects approach to combine correlation coefficients across studies in order to estimate the strength, spread, and significance of the population-level correlation. The random-effects approach reflects the fact that the studies in this meta-analysis used a variety of tasks spanning different sensory modalities and IAF calculations. Thus, rather than assuming the studies all measure the same fixed effect in the population, we assume that each experiment reflects a sample from a distribution of possible effects. This amounts to a more conservative approach in which the estimate of variance on the effect size is typically larger than in a fixed-effect meta-analysis (Field, 2001; Field & Gillett, 2010). Specifically, we used the widely-adopted random-effects approach introduced by Hunter and Schmidt (Hunter & Schmidt, 1990; Schmidt & Hunter, 2015), which amounts to a sample-size weighted average correlation and standard error (SE), which can then be used to compute a z-statistic and an associated p-value. We specifically implemented equations 11-15 as described in Field (2001) who tested the Hunter and Schmidt method against other meta-analytic estimators using simulations and found that it produced the most accurate estimate of the true population effect size and controlled for the Type 1 error rate so long as the number of studies was > 15. In addition to computing the population correlation coefficient, SE, and z-statistic, we also implemented a non-parametric bootstrap analysis (with 20,000 samples) to generate distributions of the population-level effect and to ensure that a few studies were not driving any effects observed under parametric assumptions.

We first analyzed the 27 correlations derived by averaging Fisher’s z-transformed r-values across each unique experiment. The distribution of these r-values can be seen in Figure 2A. When viewed this way it is notable that, despite several non-significant correlations in the literature, all experiments we found show an effect in the hypothesized direction with correlations ranging from a min of 0.09 to a max of 0.85. The Chi-squared test of homogeneity was non-significant (*X^2^*=32.36, p = 0.22), indicating that the distribution of correlations was approximately homogeneous and that it is appropriate to combine studies.

**Figure 2.**
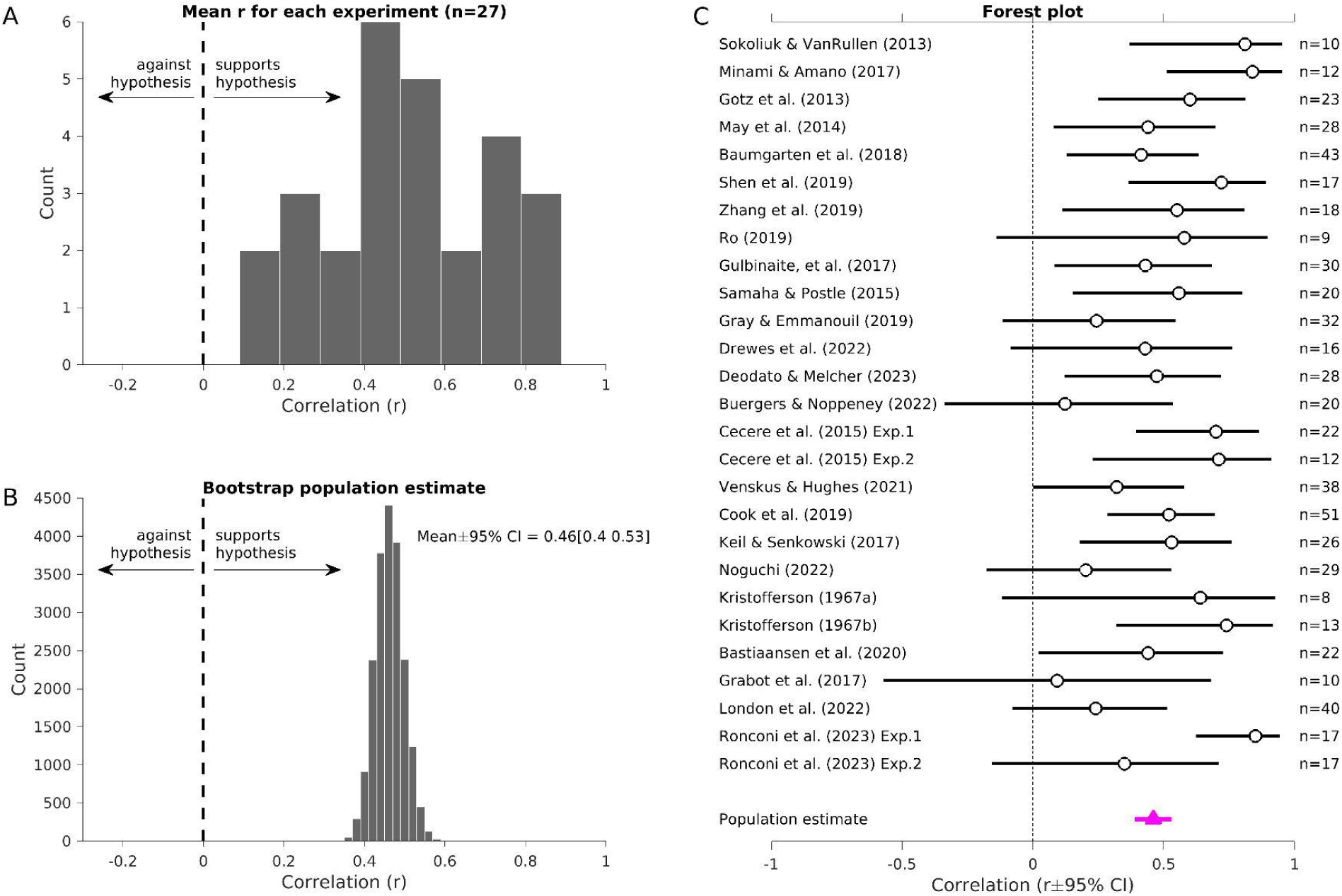
Meta-analysis of 27 experiments linking IAF to temporal properties of visual and audio-visual perception. **A)** Histogram of the mean r-values across the 27 experiments with the sign of each correlation corrected so that positive correlations support the hypothesis and negative correlations go against the hypothesis that higher IAF predicts shorter temporal windows. **B)** Bootstrap parameter distribution obtained by computing the meta-analytic r-value on 20,000 random samples (with replacement) of the 27 experiments. The distribution is tightly clustered around a population parameter estimate of r = 0.46 and does not overlap with zero. **C)** Forest plot showing the mean r-value, 95% CI, and sample size for each experiment along with the population estimate from the meta-analysis.

The meta-analysis revealed a population correlation of r(SE) = 0.460 (0.0359), suggesting a medium effect in the hypothesized direction. The tight SE obtained by combining studies results in a very large standardized effect (z-statistic) of 12.81, which was highly significant (p<0.00001). Bootstrap sampling of the population correlation effect confirmed this result with a mean r = 0.462 and 95% bootstrap confidence interval (CI) clustered between [0.393, 0.533] and non-overlapping zero. The bootstrap parameter distribution is shown in Figure 2B and a Forest plot of the r and 95% CI from each experiment and the meta-analysis is shown in Figure 2C. These results indicate that, combining across studies in the literature, there is a highly significant correlation between IAF and temporal properties of visual and audio-visual perception, with the true effect likely in the medium-to-large range between 0.39 and 0.53.

Next, we explored the effect of several experimental variables on the meta-analytic correlation estimate. Namely, we looked at whether modality (visual versus auditory-visual) and IAF derivation (eyes-open versus eyes closed) impacts the correlation strength. Analyzing only visual experiments, we again found a significant population effect (r(SE) = 0.473 (0.044), z = 10.69, p < 0.00001, bootstrap 95% CI = [0.395, 0.564]). Using only audio-visual experiments we also see a significant effect (r(SE) = 0.431 (0.056), z = 7.61, p < 0.00001, bootstrap 95% CI = [0.327, 0.547]). Although the correlation strength was numerically higher in the visual studies, the difference between bootstrapped distributions was not significant (p = 0.267). Whether a study derived IAF from eyes-open or eyes-closed data also did not have a large impact on the resulting population effect. For eyes-open IAF we found a significant meta-analytic correlation (r(SE) = 0.428 (0.038), z = 11.19, p < 0.00001, bootstrap 95% CI = [0.391, 0.505]) and the same was true for eyes-closed IAF (r(SE) = 0.474 (0.068), z = 6.93, p < 0.0001, bootstrap 95% CI = [0.305, 0.608]), although the comparatively fewer number of studies reporting eyes-closed results lead to a larger SE and therefore a smaller z-statistic.

In the above analyses, we first averaged over all the r-values reported in a single experiment to 1) get a better estimate of the true correlation in that experiment 2) only include independent experiments in our meta-analysis, and 3) and to avoid over-representing experiments that simply report more r-values (for instance, by reporting correlations with IAF from different electrodes from within the same sample of subjects). To test the influence that this analysis choice has on the results of the meta-analysis, we re-ran the analysis using all 75 correlations reported in Table 1. Note that this approach strongly weighs those studies that reported null effects. For instance, 27 out of 75 (36%) of the r-values we collected were from Buergers & Noppeney (2022) even though their experiment only contained a single sample of 20 participants. Nevertheless, even under conditions biased towards null effects, we found a highly significant meta-correlation when pooling all 75 correlations (r(SE) = 0.307 (0.028), z = 10.95, p < 0.00001, bootstrap 95% CI = [0.255 0.360]).

One possible concern is that the studies used in the present meta-analysis are subject to various forms of bias which lead to inflation of effect size estimates. One source of bias could come from making multiple statistical comparisons (e.g., across all sensors) and reporting the correlation from the largest sensor. To address this we re-ran the meta-analysis excluding four studies that performed multiple comparisons across electrodes (Cecere et al., 2015 Exp.1; Deodato & Melcher, 2023; Noguchi, 2022; Samaha & Postle, 2015). The results of this analysis remained virtually unchanged (r(SE) = 0.46 (0.039), z = 11.748, p < 0.00001, bootstrap 95% CI = [0.388 0.540]), suggesting that this type of bias had little impact on our conclusions. Additionally, as a means of standardizing the spatial selection of correlations, we ran an analysis including only those correlations which used IAF derived from occipital electrode Oz (since this was the most common electrode). Although only 7 studies (see Table 1) were included in this analysis, the effect size marginally increased and remained highly significant (r(SE) = 0.50 (0.065), z = 7.706 p < 0.0001, bootstrap 95% CI = [0.365 0.622]).

Another source of bias is the selective publication of large or significant effects while small or null effects remain unpublished. Though difficult to diagnose, a so-called “funnel plot” can be informative. As shown in Figure 3, a funnel plot displays the effect size on the x-axis (here, the correlation) by a measure of the precision of the study (here the SE of each effect on a reverse axis as recommended by Sterne et al., 2011). In the absence of bias, higher precision studies (lower SE) are expected to provide more concentrated estimates of the true effect and “funnel” out in a symmetrical way as the precision decreases (SE increases). The funnel plots in Figure 3 appear mostly symmetrical although there are a few studies that appear to have larger effect sizes than may be expected (upper right), suggesting the potential for publication bias. However, there is a general positive association between the SE and effect size such that higher precision studies tend to observe larger effects, which runs counter to the idea that spurious correlations driven by low-powered studies happen to be published and are driving the meta-analytic effects we observed. Finally, we computed Orwin’s fail-safe N (Orwin, 1983) to estimate how many unpublished studies averaging no correlation (r = 0) would need to exist in order to reduce our observed correlation to a trivial level. We adopted a standardized effect size (Cohen’s d) of 0.1 as a ‘trivial’ effect (following Hyde & Linn, 2006) and computed the fail-safe N as *N_fs_ = N_0_*((d_0_-d_c_)/d_c_)*, where N_0_ is the current number of studies (27), d_0_ is the the present effect size (Cohen’s d = 1.04, corresponding to an r of 0.46), and d_c_ is the value considered to be a negligible effect (Cohen’s d = 0.1). By this measure, it would take 252 unpublished studies averaging no effect to reduce the present meta-effect to a negligible size. Adopting a more conservative Cohen’s d of 0.2 as “negligible”, there would still need to be 112 unpublished null effects to reduce the present finding to a negligible level. Thus, while there may be some effect of publication bias it seems unlikely to be a primary driver of our results.

**Figure 3.**
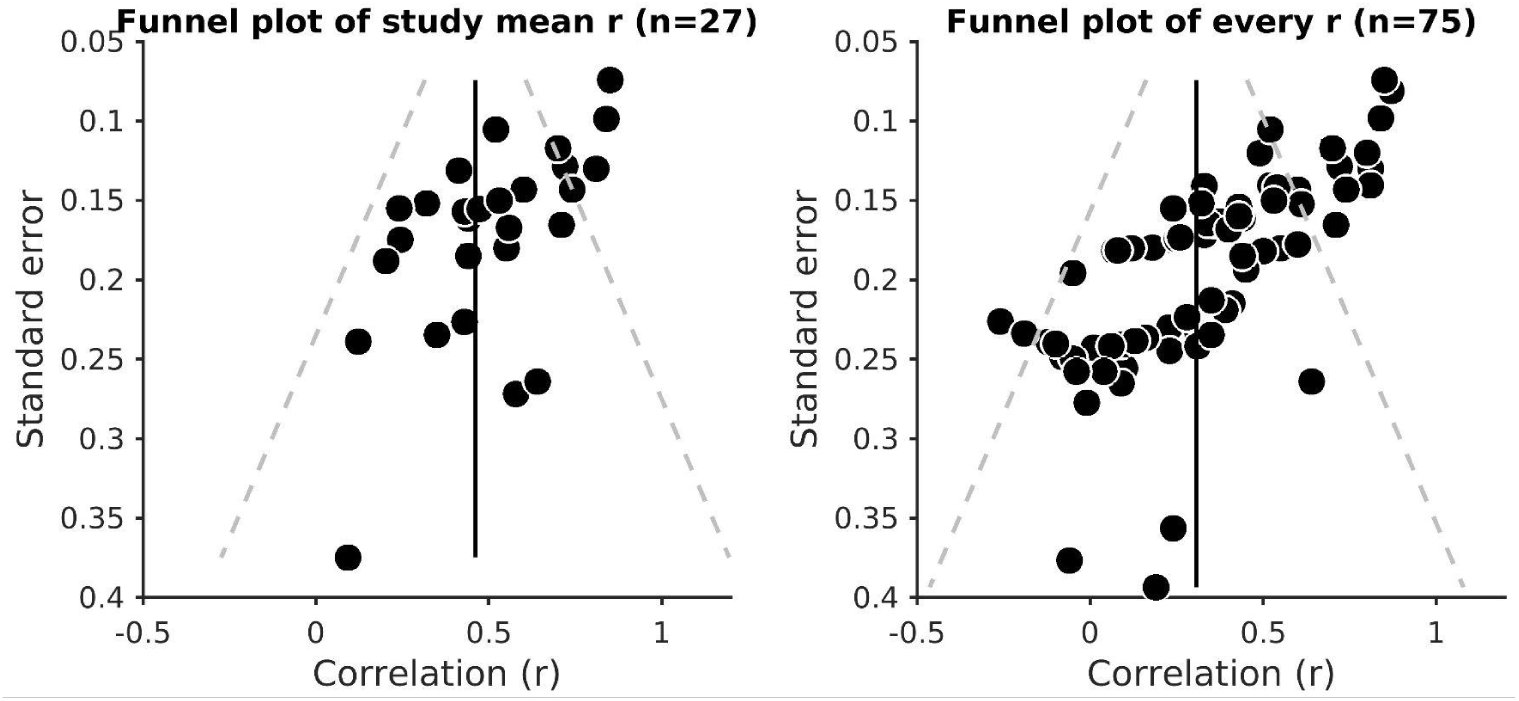
Funnel plots showing the relationship between effect size (on the x-axis) and study precision (on the y-axis; SE plotted on reverse scale). Dashed lines denote 95% confidence intervals at each SE, within which the majority of studies should fall. There are a few unexpectedly large effect size studies, which could indicate some publication bias. On the other hand, the positive association between effect size and precision suggests that more precise studies tend to find higher correlations. See text for additional analyses.

Taking these analyses together, the best population estimate suggests a correlation in the range of 0.39 to 0.53. Standard power calculations indicate that future studies should therefore aim to sample between 65 and 33 participants to achieve 90% power for a two-tailed effect or between 54 and 28 if the hypothesis justifies a one-tailed test. In sum we find strong evidence that IAF shows a medium-to-large correlation with the temporal properties of visual and audio-visual perception and that this holds for both sensory modalities, varying derivations of IAF, and even under conditions that overweights null effects in the meta-analysis.

## Comments on Buergers & Noppeney (2022)

Given the evidence for a positive effect revealed in the above meta-analysis, a question arises about how to interpret the recent null effects of Buergers & Noppeney (2022; henceforth referred to as B&N). On the one hand, given the relatively homogeneous distribution of effects we observed across studies, the null reports in the literature may simply be a matter of sampling error naturally expected on the basis of the relatively small sample sizes used in many of the experiments we looked at. That is, B&N could have been underpowered in their individual-differences analysis or just failed to observe an effect due to chance. Consistent with this idea, we note that the majority of correlations reported in B&N were actually in the predicted direction and some of them even successfully replicated (or nearly replicated) the findings of Samaha & Postle (2015), (see Table 1). An additional explanation is that perhaps particular aspects of their study design are relevant for understanding their reduced effects.

There are several differences in the task design used in B&N which B&N themselves highlight as possible sources of discrepancy. First, their task was designed to tease apart perceptual sensitivity (d’) from bias (criteria) in both visual-only 2FF and 2FF in the presence of 1 and 2 sounds. They accomplished this by always including both one and two-flash trials, whereas many studies only included a single flash (paired with two beeps in the case of the SIFI) or included only two flashes with a variable ISI (in some variants of the 2FF). We strongly endorse the use of modeling frameworks such as SDT to gain deeper insight into how neural oscillations impact perception (Di Gregorio et al., 2022; Samaha et al., 2020; Tarasi et al., 2022, 2023) and we are grateful that B&N are highlighting this issue in the field. However, we think the use of a task design tailored towards a SDT analysis is unlikely to explain the discrepant results. For one, early work in the visual domain had already used a SDT framework and found a positive effect of higher alpha frequency (within-subjects) on sensitivity in the 2FF task (Coffin & Ganz, 1977) and the effect in Samaha & Postle (2015) was obtained on 2FF discrimination thresholds (% correct) in a task with equal number of one and two-flash trials. Thus, the inclusion of one-flash trials in the visual 2FF paradigm seems unlikely to explain much. Moreover, the inclusion of two-flash trials in the multisensory context (i.e., in the SIFI) is also unlikely to be a major source of discrepancy since a recent paper used a similar stimulus configuration in a larger sample than B&N and found a correlation with IAF and the SIFI, which the author showed largely corresponded to a change in sensitivity rather than criterion (Noguchi, 2022, 2023).

A second explanation offered by B&N is that many prior studies have used arbitrary criteria for determining if a participant should be excluded from analysis on the basis of a “poorly” fitting psychometric function or “insufficient” illusion rates in the SIFI. Non-standardized rejection criteria could certainly contribute to variance in effect sizes and also introduce researcher degrees of freedom which could bias results. We again agree with B&N that these decisions should be made using objective criteria (such as those advocated for in psychophysical studies; Wichmann & Hill, 2001) and, especially where replication is concerned, be applied as consistently as possible. However, it is possible to find studies that were not subject to this issue and that still observed effects. The study by Noguchi (2022) used a single ISI and measured illusion rates and d’ at that one ISI, without fitting any psychometric functions and without excluding any participants. The study by Samaha & Postle (2015) verified their results using % correct at a fixed, 30 ms ISI, which also does not depend on psychometric function fitting. Moreover, a recent study highlights that just because a psychometric function fits the data well, this may not imply that the subject provided “good” data. Specifically, Deodato & Melcher (2023) examined correlations between IAF and 2FF in a sample of 38 participants and found that including the slope of the psychometric function as a weighting factor in the correlation greatly improved predictions. The implication is that subjects with excessively shallow slopes, despite having well-fit psychometric functions, indicate the presence of large sensory- or decision-related noise and may lead to mis-estimation (or at least added variance in the estimation) of 2FF thresholds.

We offer some additional considerations for the reduced effects reported in B&N. The foremost being that the size and eccentricity of the stimuli used (a 1.34° SD Gaussian light blob presented at 15° leftward eccentricity) places the visual stimulus (presumably unintentionally) squarely in the average human blind spot (Safran et al., 1993). A potential consequence is that the task is likely being performed monocularly for some subjects but not others, given natural variation in location of each person’s blindspot. It has previously been shown that 2FF thresholds are higher (performance is worse) during monocular compared to binocular viewing (Pearson & Tong, 1968). Importantly, it has also been found that the SIFI is reduced and even sometimes abolished under monocular viewing conditions (Moro & Steeves, 2018). Thus, it is likely that the results of B&N are affected by an additional source of between-subjects variance (the extent to which some participants were doing the task monocularly versus binocularly) since it is known that monocular viewing impacts both 2FF thresholds and the SIFI.

Additionally, the stimuli used in Samaha & Postle (2015) were designed to rule out total stimulus duration as a possible confounding cue observers could use to make their one versus two flash judgments since manipulating only the ISI on two-flash trials will produce an overall longer stimulus sequence. The direct replication of the Samaha & Postle stimulus parameters in Gray & Emmanouil (2020) led to a successful replication of the correlation with IAF in a sample of 31 subjects. In contrast, B&N used a range of ISIs from 23 to 223 ms but did not match the duration of one-flash trials. Given that subjects performed 5 days of testing and that the JND for duration of very brief visual events is around 30 ms (Rammsayer et al., 2015), the well-trained subjects in B&N could have learned to incorporate overall stimulus duration as a cue, rather than solely relying on the number of flashes perceived.

Lastly, a close examination of the reliability of some of the behavioral data in B&N could be useful, particularly with respect to the SIFI conditions. In their condition which most closely approximates the design of Cecere et al. (2015) and Cooke et al. (2019), which B&N refer to in their paper as “yes-no” SOA, two flashes are presented along with one or two sounds. Their Figure 4 shows that 7 out of the 19 subjects included in that analysis have thresholds at the largest ISI used in their experiment (223 ms), suggesting these participants have exceedingly large binding windows for the SIFI that were not captured well by the stimulus range or they were performing the task using some different strategy. The remaining participants in the scatter plot, whose thresholds are in the range observed in the other SIFI experiments reviewed here (between 40 and 140 ms) appear to show the negative trend expected as a function of IAF. Consistent with this worry about threshold reliability, neither of the SIFI thresholds estimated in the other tasks (2IFC or staircase procedure) produced strong between-task correlations (see B&N supplementary Figure 7). In other words, the SIFI threshold estimated via psychometric function fitting were uncorrelated with the SIFI threshold estimated via staircase and neither were strongly correlated with the threshold estimates via 2IFC. This implies that the implementation of the measurement in that experiment may not have been very reliable, which has the well-known consequence of limiting the extent to which the measure can correlate with other measures, such as IAF (Spearman, 1904).

## Open questions and future directions

We conducted a meta-analysis of 27 studies that investigated the correlation between IAF and various temporal properties of visual and audio-visual perception. Our results estimate the true correlation value in the population to be around 0.39 to 0.53, a medium-to-large effect. Similar effects were found when considering only visual or only audio-visual temporal binding windows in isolation as well as when examining different IAF estimation protocols (i.e., eyes-open versus eyes-closed). Moreover, using analysis choices that favor null results, we still find a significant population effect. Our meta-analysis points to an effect in the predicted direction, however a number of open questions remain and several recommendations for future research can be made.

Chief among them is, what are the mechanisms underlying the link observed in the correlational studies reviewed here? A recent distinction has been introduced between *rhythmic* and *discrete* perception (VanRullen, 2016) whereby alpha oscillations either phasically modulate the excitability of neural processes and bias the strength of perceptual experiences (rhythmic account) or they shift the timing of neural events such that the perceived order of stimuli is alpha-phase dependent. Recent work has found that ongoing alpha phase largely modulates the amplitude of early visual responses rather than their latency (Dou et al., 2022) and suggests that IAF is not predictive of individual differences in the latency of early visual ERP components (Morrow et al., 2023), both results that could be seen to favor rhythmic over discrete perception. From this perspective, how could the effects of IAF on the temporal properties of perception reviewed here be explained? In the audio-visual domain, alpha has been proposed to reflect that latency of audio-visual interactions, such that the first beep in the SIFI, for instance, has an excitatory influence on visual cortex at a delay specified by the alpha period, possibly via phase-reset mechanisms (Mercier et al., 2013; Romei et al., 2012). Then, when the second sound arrives, the visual cortex is in a more excitable state and the second sound input causes the illusory perception of a flash (Cecere et al., 2015). A similar excitability-based (rather than timing-based) mechanism was proposed to underlie the relationship between IAF and the triple flash illusion (Gulbinaite et al., 2017). The distinction between rhythmic and discrete effects of alpha, we think, should motivate the use of new perceptual tasks that better delineate between excitability versus timing effects. For instance, although the 2FF paradigm was widely used across the papers we reviewed, is it hard to distinguish whether alpha frequency is influencing the binding (and hence, probably the perceived timing) of the two flashes, or whether one flash is simply more likely to be missed, causing the percept of a single flash (Fan, 2018; Samaha & Postle, 2015). In other words, tasks that more directly assess temporal binding windows as opposed to just temporal *influence* windows should be considered (Wutz et al., 2018).

We also have hope that model-based approaches to measuring relevant perceptual phenomena hold promise in helping to elucidate underlying mechanisms and guide experimental design. Not many of the reviewed studies have used tasks that allow for distinguishing between sensitivity and criterion effects. Those that have either found null effects of IAF on both (Buergers & Noppeney, 2022), or effects on sensitivity and not criterion (Coffin & Ganz, 1977; Noguchi, 2022, 2023). This also leads into a question about how best to estimate perceptual thresholds. Given that yes-no tasks and discrimination tasks can be influenced by either sensitivity or criterion shifts, methods that provide bias-free threshold estimates might be preferred. For instance, using SDT to estimate thresholds or adopting less-biased 2AFC or 2IFC approaches could be recommended. However, as Buergers & Noppeney (2022) results show, at least in the case of perceptual illusions like the SIFI, these different procedures can provide threshold estimates that bear little resemblance to one another. A further consideration is that many genuine perceptual illusions may be best characterized as criterion effects, rather than sensitivity changes (Knotts & Shams, 2016). Thinking beyond threshold measures, the deleterious effects that shallow psychometric slopes can have on IAF-threshold correlations (Deodato & Melcher, 2023) may also motivate a method-of-constant stimuli approach (allowing for a full psychometric function to be fit) over a simple threshold-estimation procedure (e.g., a staircase). Lastly, we wish to emphasize the sample-size recommendations that should be considered for detecting this effect in future research. Our results suggest a population correlation parameter of ∼0.46, requiring a sample size of 45 if 90% power and a two-tailed test are desired.

Our paper focused on IAF and individual differences in temporal properties of visual and audio-visual perception since this scope provided enough experiments for meta-analyses. However, relevant evidence also comes from experiments that examined within-subject (i.e., trial-to-trial) fluctuations in alpha frequency. For instance, Coffin & Ganz (1977) first noted (and replicated) that trials with higher pre-stimulus alpha frequencies were associated with greater 2-flash percepts in the 2FF paradigm. Several studies have subsequently replicated this effect (Drewes et al., 2022; Samaha & Postle, 2015; Shen et al., 2019) and have even shown that manipulating task demands to favor integration or segregation of an identical stimulus sequence can slow down or speed up alpha frequency, respectively (Han et al., 2023; Sharp et al., 2022; Wutz et al., 2018), suggesting top-down regulation of alpha frequency as a mechanism to flexibly adapt visual sampling. It is worth noting, however, that Buergers & Noppeney (2022) did not replicate these within-subject effects of alpha frequency, an effect they were considerably better powered to detect given the large number of trials in their experiment. Although there are not enough within-subject studies yet to perform the kind of meta-analysis we conducted here for between-subject effects, we can speculate that the design choices articulated in the previous section could have influenced the null within-subjects result obtained in Buergers & Noppeney (2022) as well. It is also worth noting that the common measure used to estimate within-subject frequency shifts (so-called “frequency sliding”) has recently been found to be biased by other factors such as power changes or changes in the spectrum slope (Samaha & Cohen, 2022), which means it should be applied cautiously.

Lastly, two critical yet unanswered questions concern the generality of IAFs predictive power. First, is alpha frequency a general binding mechanism across sensory modalities? And second, is alpha frequency reflective of some general timing mechanism in cognition more broadly, beyond low-level perceptual interactions. Regarding the first question, our review focused on the two modalities with the most experiments (vision and audio-visual binding), but work in the tactile domain has also led to interesting effects. Migliorati et al. (2020) found that IAF predicted the size of the temporal binding window for visuotactile stimuli using a SJ task. On the other hand the tactile equivalent of the SIFI was found to correlate with individual beta frequency, and not alpha (Cooke et al., 2019; Fotia et al., 2021). Moreover, the link between beta-band oscillations and a tactile resolution (measured using a tactile variant of the 2FF) has been mixed (Baumgarten et al., 2015, 2017). Regarding the second question, higher alpha frequency has been linked to greater perceptual sensitivity in tasks that seemingly do not depend on temporal binding, such as detection (Tarasi & Romei, 2023) and discrimination of contrast-defined targets (Di Gregorio et al., 2022; Nelli et al., 2017; Trajkovic et al., 2023), although one could argue that these tasks also benefit from more frequent sampling given the brief nature of the stimuli. More generally though, IAF has been found to correlate with measures of processing speed and performance metrics in a variety of cognitive tasks beyond perception (Angelakis et al., 2004; Bertaccini et al., 2022; Klimesch et al., 1993; Samuel et al., 2018; Surwillo, 1961). This prompts the hypothesis that alpha frequency effects on perception may be something of a byproduct of overall enhanced cognitive performance or task engagement rather than a low-level binding mechanism. Or, speculatively, the arrow of causality is reversed and the temporal constraints imposed by alpha on low-level processing serve as a building block for more complex cognitive tasks. We hope future research can shed light on these two questions.

